# CircRNA-regulated immune response of Asian honey bee workers to microsporidian infection

**DOI:** 10.1101/2022.06.30.498258

**Authors:** Zhiwei Zhu, Jie Wang, Xiaoxue Fan, Qi Long, Huazhi Chen, Yaping Ye, Kaiyao Zhang, Zhongmin Ren, Yang Zhang, Qingsheng Niu, Dafu Chen, Rui Guo

**Affiliations:** College of Animal Sciences (College of Bee Science), Fujian Agriculture and Forestry University, Fuzhou 350002, China; Apitherapy Research Institute, Fujian Agriculture and Forestry University, Fuzhou 350002, China; Apiculture Science Institute of Jilin Province, Jilin 132000, China

**Keywords:** *Apis cerana cerana*, circular RNA, *Nosema ceranae*, non-coding RNA, immune response, host-pathogen interaction

## Abstract

*Nosema ceranae* is a widespread fungal parasite for honey bees, causing bee nosemosis. Based on deep sequencing and bioinformatics, identification of circular RNAs (circRNAs) in *Apis cerana cerana* workers’ midguts and circRNA-regulated immune response of host to *N. ceranae* invasion were conducted in this current work, followed by molecular verification of back-splicing sites and expression trends of circRNAs. Here, 10185 and 7405 circRNAs were identified in the midguts of workers at 7 d (AcT1) and 10 d (AcT2) post inoculation (dpi) with *N. ceranae*. PCR amplification result verified the back-splicing sites in three specific circRNAs (novel_circ_005123, novel_circ_007177, and novel_circ_015140) expressed in *N. ceranae*-inoculated midgut. In combination with transcriptome data from corresponding un-inoculated midguts (AcCK1 and AcCK2), 2266 circRNAs were found to be shared by the aforementioned four groups, whereas the numbers of specific ones were 2618, 1917, 5691 and 3723 respectively. Further, 83 (52) differentially expressed circRNAs (DEcircRNAs) were identified in AcCK1 vs AcT1 (AcCK2 vs AcT2) comparison group. Source genes of DEcircRNAs in workers’ midgut at 7 dpi were involved in two cellular immune-related pathways such as endocytosis and ubiquitin mediated proteolysis. Additionally, competing endogenous RNA network analysis showed that 23 (13) DEcircRNAs in AcCK1 vs AcT1 (AcCK2 vs AcT2) can target 18 (14) miRNAs and further link to 1111 (1093) mRNAs. These target mRNAs were annotated to six cellular immunity pathways including endocytosis, lysosome, phagosome, ubiquitin mediated proteolysis, metabolism of xenobiotics by cytochrome P450, and insect hormone biosynthesis. Moreover, 284 (164) IRES and 54 (26) ORF were identified from DEcircRNAs in AcCK1 vs AcT1 (AcCK2 vs AcT2) comparison group; additionally, ORFs in DEcircRNAs in midgut at 7 dpi with *N. ceranae* were associated with several crucial pathways including endocytosis and ubiquitin-mediated proteolysis. Finally, RT-qPCR results showed that the expression trends of six DEcircRNAs were consistent with those in transcriptome data. These results demonstrated that *N. ceranae* altered the expression pattern of circRNAs in *A. c. cerana* workers’ midguts, and DEcircRNAs were likely to regulate host cellular and humoral immune response to microsporidian infection. Our findings lay a foundation for clarifying the mechanism underlying host immune response to *N. ceranae* infection and provide a new insight into interaction between Asian honey bee and microsporidian.

## 1. Introduction

As a novel player in the world of non-coding RNA (ncRNA), circular RNA (circRNA) has become a worldwide research hotspot. Different from canonical alternative splicing, circRNA is generated from back-splicing of pre-mRNA (Li et al., 2018). In comparison with linear RNA, circRNA is more resistant to RNase R enzyme digestion due to its special covalently closed-loop structure, hence circRNA is regarded as ideal endogenous biomarker (Meng et al., 2017). CircRNAs are abundant in eukaryotic cells and play versatile function such as regulation of source gene’s transcription (Li et al., 2015), absorption of miRNAs or RNA binding proteins as “molecular sponges” (Han et al., 2020), and translation into pepetides or proteins (Wang et al., 2020). Accumulating evidence suggest that circRNAs are involved in the occurrence and development of human diseases such as cervical (Chen et al., 2019b) and lung cancers (Wang et al., 2020). The biological function of circRNAs as competing endogenous RNAs (ceRNAs) has only been deeply studied in human and few other model species (Li et al., 2020; Han et al., 2020). For example, Li et al. reported that circTLK1 was highly expressed in renal cell carcinoma (RCC) and could promote RCC progression through the miR-136-5p/CBX4 pathway, and circTLK1 could serve as diagnostic molecule and therapeutic target for renal cancer (Li et al., 2020). However, study on insect circRNAs was very lagging and limited work was mainly focused on *Drosophila melanogaster* (Westholm et al., 2014; Krishnamoorthy A and Kadener S, 2021; Huang C et al., 2018), *Bombyx mori* (Gan et al., 2017; Hu et al., 2018a; Hu et al., 2018b; Wang et al., 2015), and honey bee (Chen et al., 2019c; Thölken C et al., 2019; Chen et al., 2020a). Westholm’s group predicted more than 2500 circRNAs using transcriptome data from *Drosophila*, and revealed circRNAs were abundantly expressed in the brain and accumulated over time (Westholm et al., 2014). Wang et al. discovered that *Bombyx mori* CircEgg was mainly located in the cytoplasm and circEgg overexpression inhibited the production of linear transcripts of *BmEgg*, circEgg can inhibit methylation of histone H3 lysine 9 by acting as the “sponge” of bmo-miR-3391-5p (Wang et al., 2020). Our group previously conducted comprehensive investigation of circRNAs in the midguts of European honey bee and Asian honey bee, Chen et al. identified 1101 circRNAs in *Apis mellifera ligustica* workers using a combination of RNA-seq and bioformatics (Chen et al., 2020a), Xiong et al. analyzed the expression profile of circRNAs in the developmental process of *Apis cerana cerana* workers’ midguts, and revealed the potential regulatory role of differentially expressed circRNAs (DEcircRNAs) (Xiong et al., 2018).

*Nosema ceranae*, an emergent fungal parasite, specifically infects the midgut epithelial cells of bee host (Huang and Evans, 2020). It was first identified in *Apis cerana* colonies reared in China (Fries et al., 1996), and then spread to *A. mellifera* colonies in Europe (Higes et al., 2010), America (Emsen et al., 2020), and other parts of the world (Giersch et al., 2009). *N. ceranae* spores enter the midgut of bee host through ingestion of contaminated food or water and then germinate due to activation by the special physical and chemical conditions inside the midgut. The infective sporoplasm is injected into the host midgut epithelium and replicated by stealing host material and energy. With the increase of the quantity of spores, the host cell finally ruptured and the released spores in the feces become new sources of infection via feeding and cleaning activities inside the colonies, or are disseminated into the environment (Higes et al., 2007; Gisder et al., 2011). *N. ceranae* infection has negative influence on bee host, such as damage of midgut epithelial cells, energy stress, immunosuppression, cell apoptosis inhibition, and lifespan reduction (Goblirsch et al., 2013; Antúnez et al., 2009; Mayack and Naug, 2009; Kurze et al., 2018; Panek et al., 2018). Additionally, *N. ceranae* can severely weaken bee colony health and productivity in collaboration with other biological or environmental stress (Doublet et al., 2015). *A. c. cerana*, a subspecies of *A. cerana*, is mainly distributed and widely used in Asian countries including China. Compared with western honey bee, *A. cerana* has several obvious advantages, e.g. *A. cerana* is more adaptive to extreme weather conditions and good at collecting scattered nectar sources (Zhao et al., 2020). Additionally, *A. c. cerana* has been used as a model for investigating of host-pathogen interaction (Huang et al., 2018). The reference genome of *A. cerana* (Park et al., 2015) was published in 2015, much later than the *A. mellifera* genome (Honeybee Genome Sequencing Consortium, 2006). Currently, study on omics and molecular biology of *A. cerana* is lagging when compared with *A. mellifera*, and interaction between *A. cerana* and parasites or pathogens is still largely unknown. Our team previously investigated the immune response of *A. c. cerana* workers to *N. ceranae* infection (Xing et al., 2021), and deciphered the differential expression profile of host miRNAs during microsporidian infection and DEmiRNA-regulated host response (Chen et al., 2019a).

CircRNAs had been identified in both *A. c. cerana* (Chen et al., 2020a) and *N. ceranae* (Guo et al., 2018b) by our group. CircRNAs have been suggested to crucial regulators involved in host-pathogen interaction. However, study on interaction between Asian honey bee and *N. ceranae* is still lacking until now. Our group previously conducted deep sequencing of *A. c. ceranae* workers’ midgut tissues at 7 days post inoculation (dpi) and 10 dpi with *N. ceranae* (AcT1 and AcT2 groups) and corresponding un-inoculated midgut tissues (AcCK1 and AcCK2 groups), and identified 9589 circRNAs based on transcriptome data from control groups (Chen et al., 2020a). Here, in order to uncover circRNA-regulated responses of Asian honey bee workers to *N. ceranae* infection, the differential expression pattern of circRNAs in *A. c. cerana* workers’ midguts responding to *N. ceranae* invasion was analyzed utilizing the obtained high-quality transcriptome data, followed by in-depth investigation of host response mediated by DEcircRNAs, with a focus on cellular and humoral immune responses. To the best of our knowledge, this is the first documentation of circRNA-regulated response of bee host to microsporidian infection. Findings in this current work can not only lay a key foundation for clarifying the underlying mechanism but also provide a novel insight into Asian honey bee-microsporidian interaction.

## 2. Results

### 2.1 Quality control of transcriptome data from *N. ceranae*-inoculated groups

Based on strand-specific cDNA library-based transcriptome sequencing, 524100096 and 615893838 raw reads were generated from AcT1 and AcT2 groups, and 515604182 and 601712328 clean reads were gained after quality control, respectively; additionally, 338500740 and 570789154 anchors reads were identified, among which 31889778 (28.28%) and 35037206 (20.60%) were aligned to the reference genome of *A. cerana*.

### 2.2 Identification, analysis, and validation of circRNAs in *N. ceranae*-inoculated *A. c. ceranae* workers’s midguts

In total, 10185 and 7405 circRNAs were identified in AcT1 and AcT2 groups, respectively. Combined with circRNAs previously identified in un-inoculated groups, Venn analysis indicated that 2266 circRNAs were shared by AcCK1, AcCK2, AcT1, and AcT2 groups, whereas the numbers of specific circRNAs were 2618, 1917, 5691, and 3723, respectively (**Fig. 1a**). Additionally, annotated exonic circRNA was the most abundant type in both AcT1 and AcT2 groups, followed by antisense circRNA and single exonic circRNA (**Fig. 1b**). Moreover, the length of circRNAs in *N. ceranae*-inoculated groups was ranged from 1 nt to more than 2000 nt, and circRNAs with the length distribution among 401-600 nt were the most (**Fig. 1c**).

**Fig. 1.**
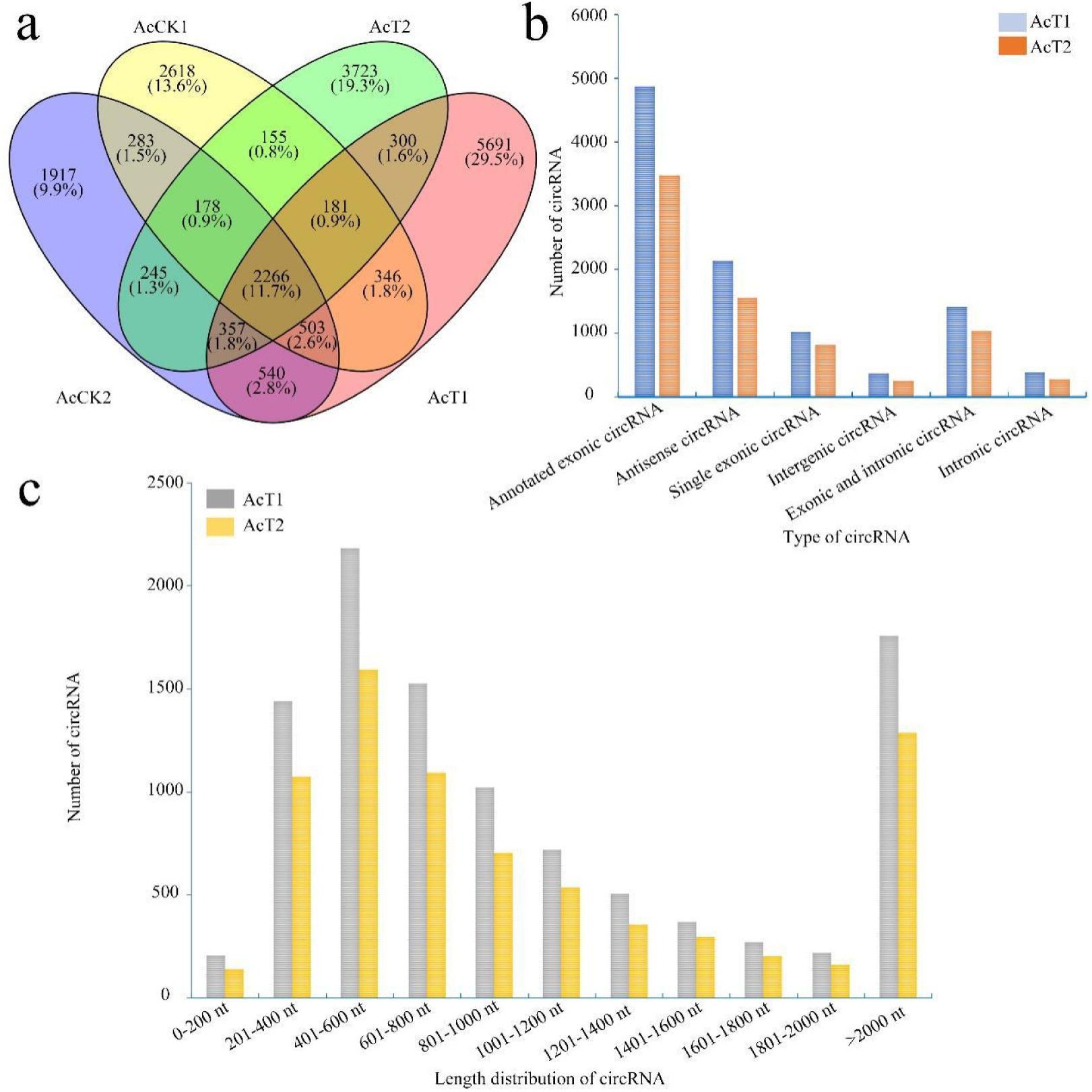
Number, type, and length distribution of circRNAs in the midguts of *A. c. cerana* workers inoculated with *N. ceranae*. (a) Venn diagram of circRNAs in AcCK1, AcCK2, AcT1, and AcT2 groups. (b) Number statistics of circRNAs derived from various genomic origins. (c) Length distribution of circRNAs.

PCR amplification was performed to further validate the three specific circRNAs identified in *N. ceranae*-inoculated midguts, and the AGE suggested that the fragments with expected sizes could be amplified using specific divergent primers for novel_circ_005123, novel_circ_007177, and novel_circ_015140 (**Fig. 2a**). Additionally, the back-splicing sites of selected circRNAs were successfully detected using Sanger sequencing (**Fig. 2b**).

**Fig. 2.**
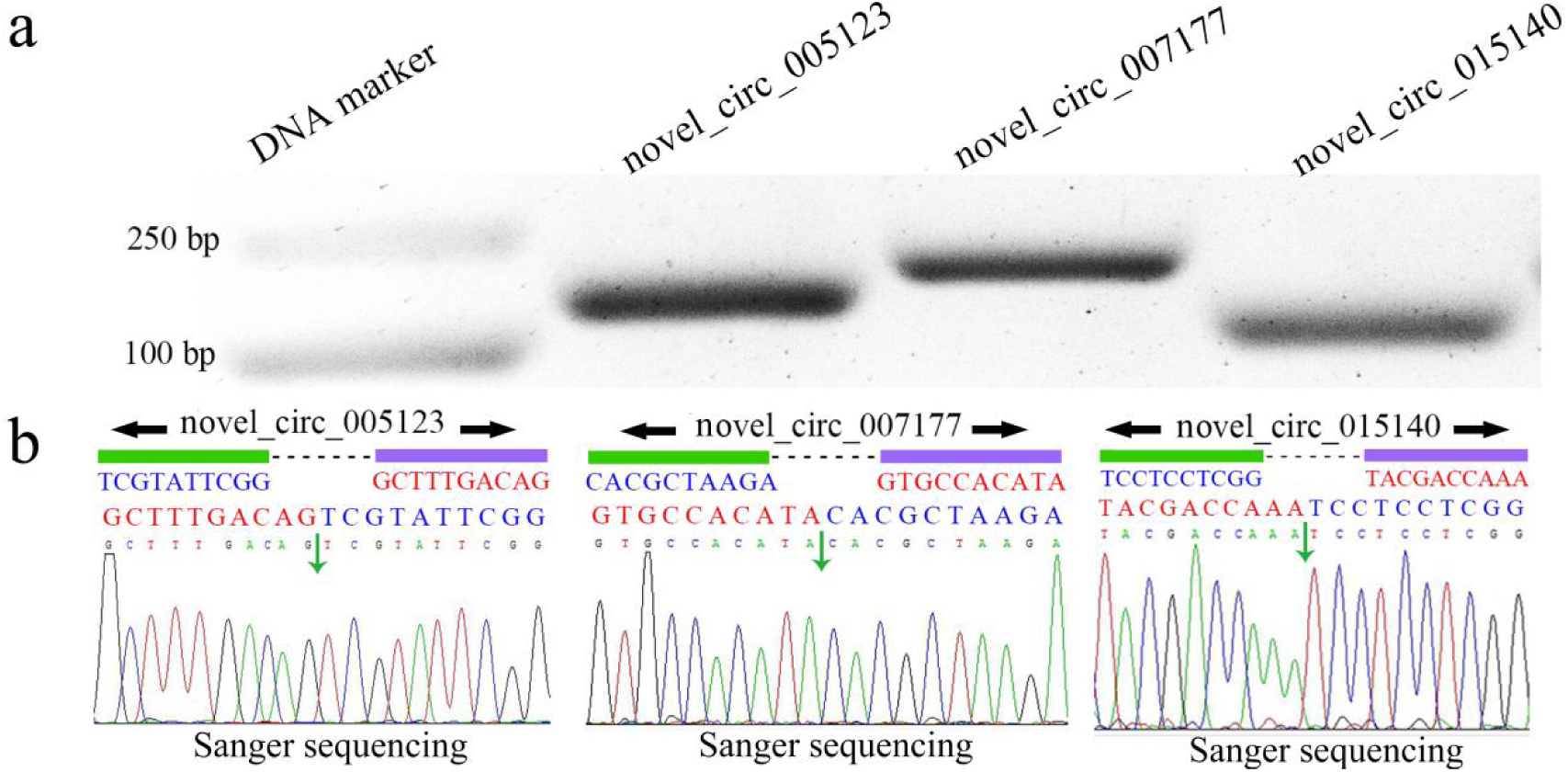
PCR amplification (a) and Sanger sequencing (b) of three *A. c. cerana* circRNAs. Black arrows indicate the direction of primers for PCR amplification, and green arrows indicate back-splicing sites of circRNAs.

### 2.3 Differentially expressed pattern of circRNAs engaged in host response to *N. ceranae* infection

A total of 83 DEcircRNAs were identified in AcCK1 vs AcT1 comparison group, including 57 up-regulated circRNAs and 26 down-regulated ones **(Fig. 3a, see also Supplementary Table S1)**; the expression levels of DEcircRNAs were among 0.001∼353.49; the most up-regulated and down-regulated circRNAs were novel_circ_012754 (log_2_FC=18.16) and novel_circ_017486 (log_2_FC=-17.05), respectively. In AcCK2 vs AcT2 comparison group, 52 DEcircRNAs were identified, including 28 up-regulated circRNAs and 24 down-regulated ones **(Fig. 3a, see also Supplementary Table S2)**; the expression levels of circRNAs were among 0.001∼445.77; the most up-regulated and down regulated circRNAs were novel_circ_002265 (log_2_FC =18.77) and novel_circ_011100 (log_2_FC =-17.50), respectively. In addition, no DEcircRNA was shared by above-mentioned two comparison groups (**Fig. 3b**).

**Fig. 3.**
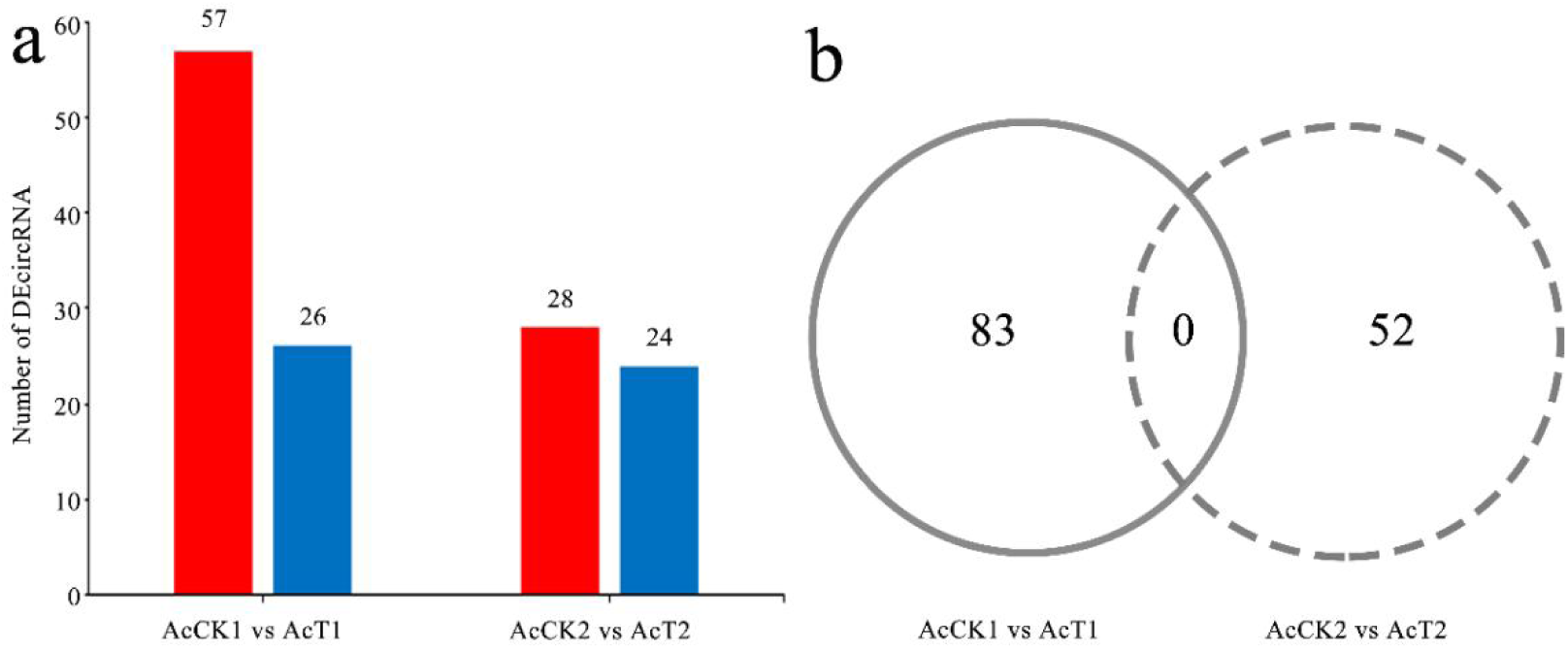
Number statistics of DEcircRNAs in AcCK1 vs AcT1 and AcCK2 vs AcT2 comparison groups. (a) Number of up- and down-regulated circRNAs (b) Venn analysis of DEcircRNAs

### 2.4 GO term and KEGG pathway analyses of source genes of host DEcircRNAs

GO classification suggested that 78 source genes DEcircRNAs in AcCK1 vs AcT1 comparison group were predicted; among these, 23 source genes were classified into 10 functional terms associated with molecular function, cellular component, and biological process, such as binding, localization, and membrane (**Fig. 4a**). Additionally, 45 sources genes of DEcircRNAs in AcCK2 vs AcT2 comparison group were predicted, among which 13 were grouped into 10 functional terms including catalytic activity, cell part, and metabolic processes (**Fig. 4b**).

**Fig. 4.**
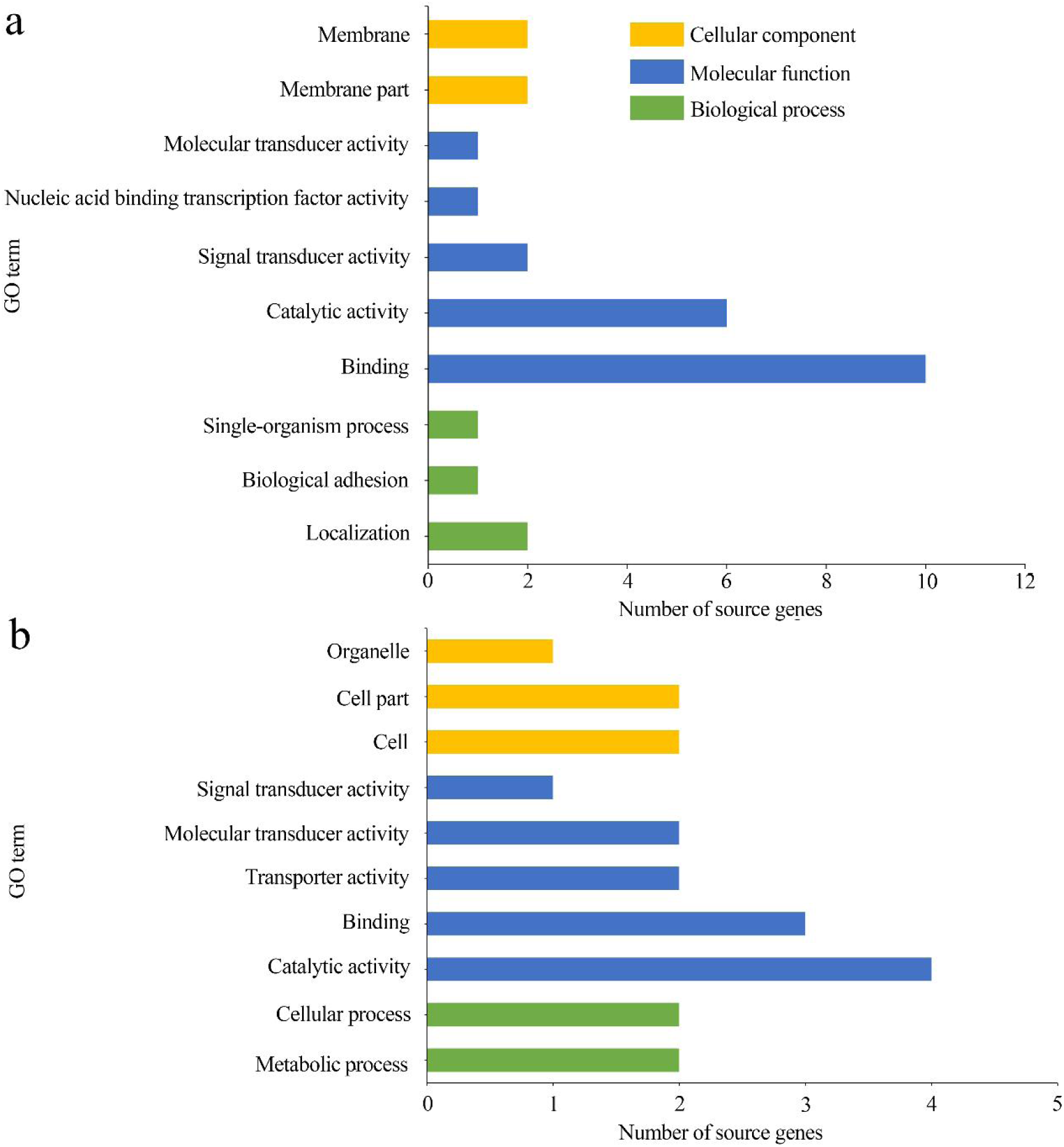
GO categorization of DEcircRNAs’ source genes in AcCK1 vs AcT1 (a) and AcCK2 vs AcT2 (b) comparison groups.

**Fig. 5.**
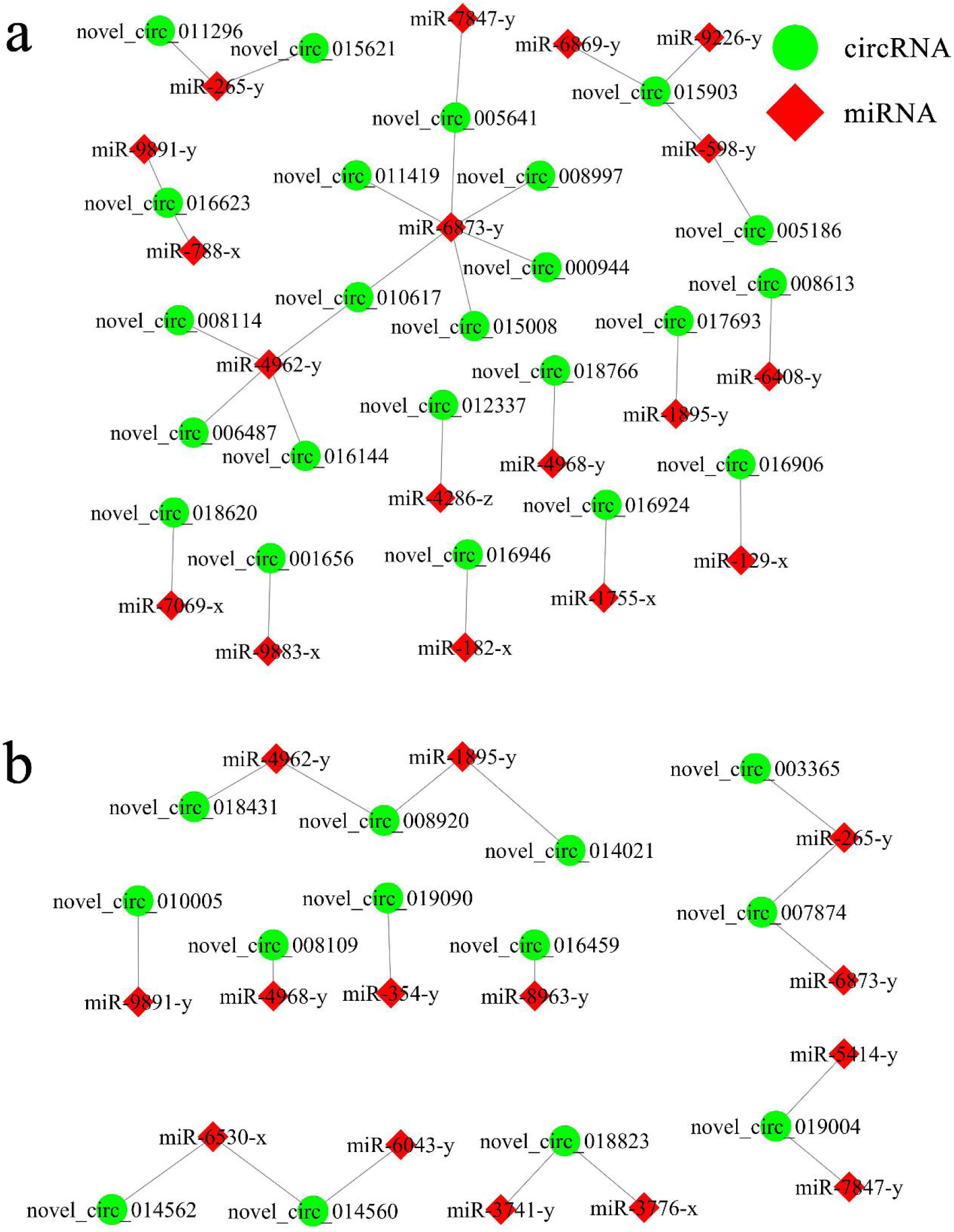
DEcircRNA-miRNA network engaged in *N. ceranae*-response of *A. c. cerana* workers. (a) Regulatory network in AcCK1 vs AcT1 comparison group (b) Regulatory network in AcCK2 vs AcT2 comparison group

**Fig. 6.**
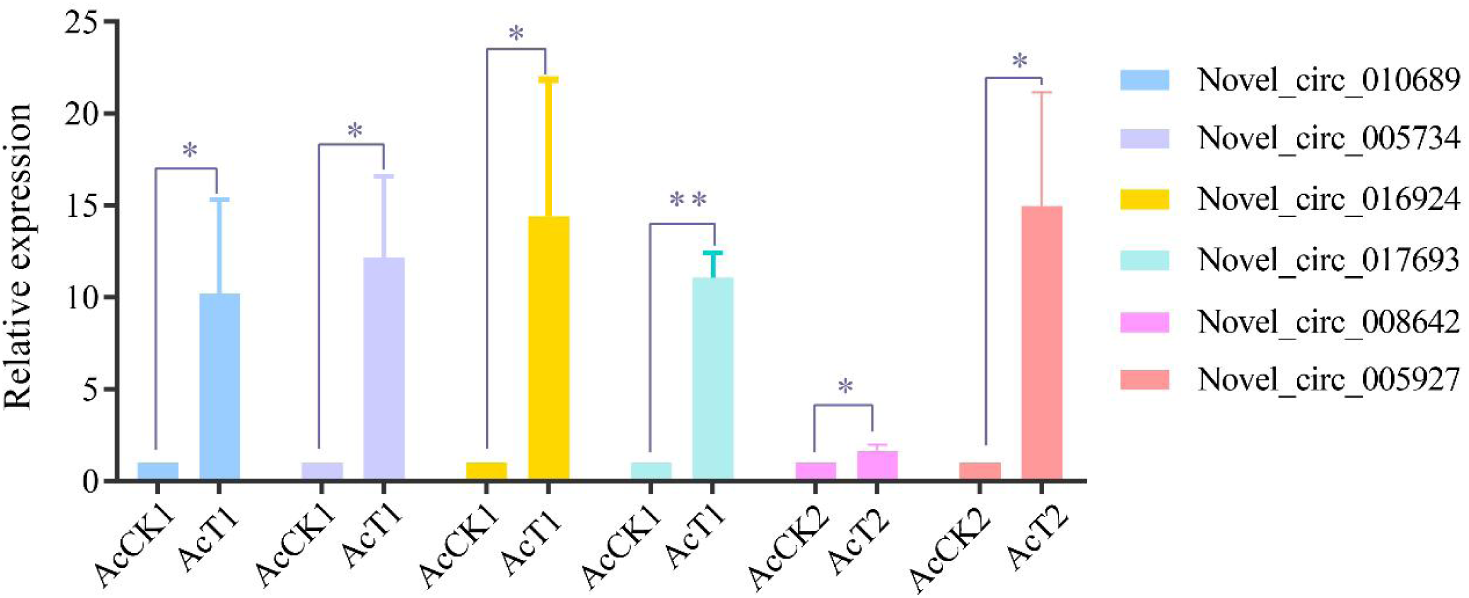
RT-qPCR verification of DEcircRNAs. Student’s *t* test, “*” indicates *P*≤0.05 and “**” indicates *P*≤0.01.

**Fig. 7.**
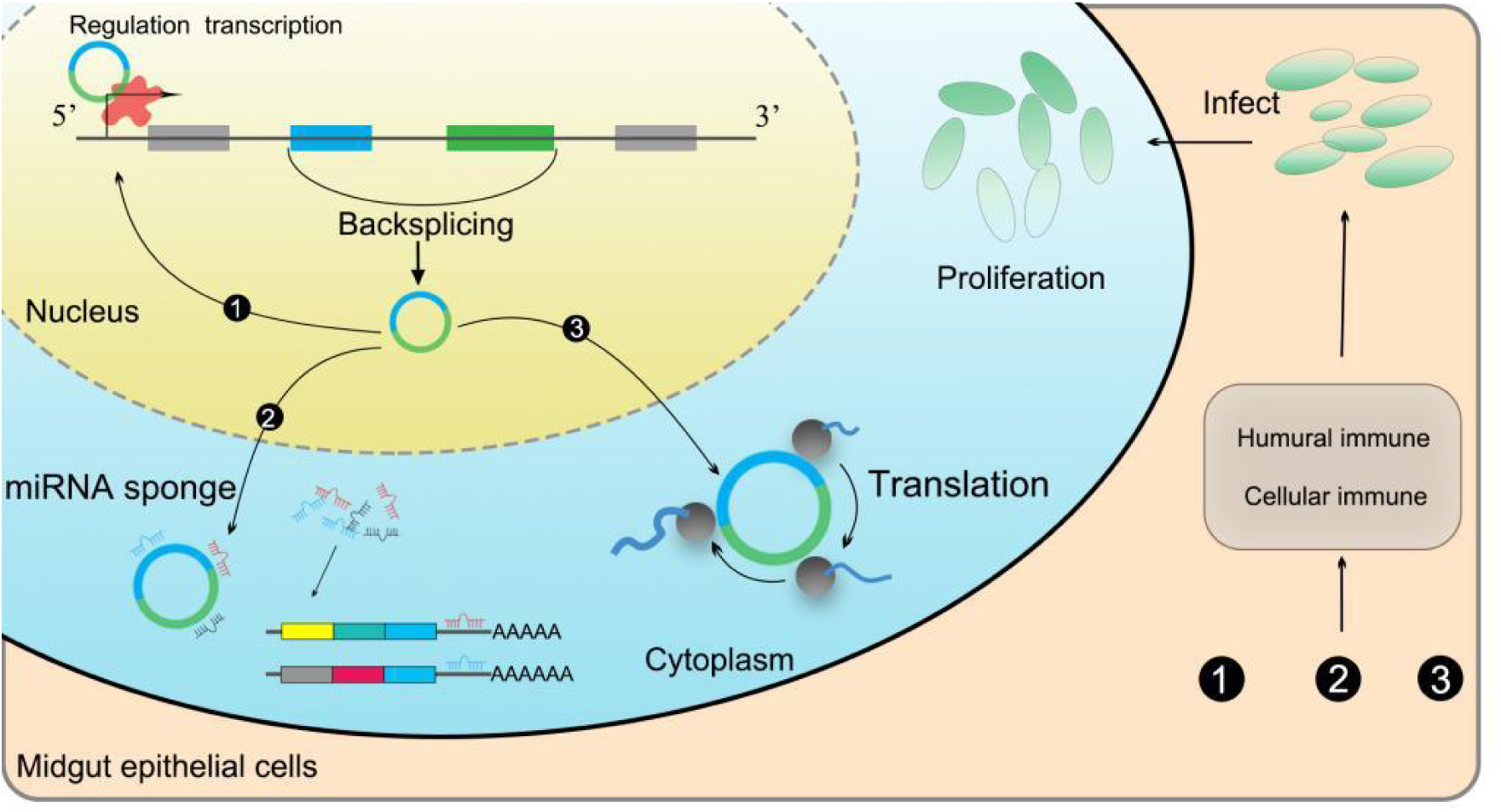
A working model of DEcircRNA-regulated response of *A. c. cerana* workers to *N. ceranae* infection.

KEGG pathway analysis suggested that source genes of DEcircRNAs in AcCK1 vs AcT1 comparison group were involved in 12 pathways such as endocytosis and Hippo signaling pathway (**Table 5**). Comparatively, source genes of DEcircRNAs in AcCK2 vs AcT2 comparison group were engaged in five pathways such as sphingolipid metabolism and mTOR signaling pathway (**Table 6**).

**Table 5.** KEGG pathways enriched by source genes of DEcircRNAs in AcCK1 vs AcT1 comparison group

| Pathway | Ko number | Number of source gene | Gene ID |
| --- | --- | --- | --- |
| Spliceosome | ko03040 | 2 | 107997744, 108004399 |
| Endocytosis | ko04144 | 2 | 107993574, 108001896 |
| Mucin type O-glycan biosynthesis | ko00512 | 1 | 108003572 |
| Tyrosine metabolism | ko00350 | 1 | 107994400 |
| Other glycan degradation | ko00511 | 1 | 108001667 |
| ECM-receptor interaction | ko04512 | 1 | 107997219 |
| Insulin resistance | ko04931 | 1 | 107999877 |
| FoxO signaling pathway | ko04068 | 1 | 107999607 |
| Hippo signaling pathway-fly | ko04391 | 1 | 107995510 |
| Ribosome biogenesis in eukaryotes | ko03008 | 1 | 107995090 |
| Ubiquitin mediated proteolysis | ko04120 | 1 | 108001789 |
| RNA transport | ko03013 | 1 | 107998561 |

**Table 6.** KEGG pathways enriched by source genes of DEcircRNAs in AcCK2 vs AcT2 comparison group

| Pathway | Ko number | Number of source gene | Source gene ID |
| --- | --- | --- | --- |
| Sphingolipid metabolism | ko00600 | 1 | 107995902 |
| ECM-receptor interaction | ko04512 | 1 | 107998342 |
| mTOR signaling pathway | ko04150 | 1 | 107997811 |
| Insulin resistance | ko04931 | 1 | 107999331 |
| Neuroactive ligand-receptor interaction | ko04080 | 1 | 107993584 |

### 2.5 DEcircRNA-miRNA regulatory network involved in *N. ceranae*-response of *A. c. cerana* workers

In AcCK1 vs AcT1 comparison group, 23 DEcircRNAs were predicted to target 18 miRNAs; among these DEcircRNAs, novel_circ_015903 could target three miRNAs, both novel_circ_016623 and novel_circ_010617 could target two miRNAs, while another 20 DEcircRNAs can target only one miRNA (**Fig. 4a**). In addition, 13 DEcircRNAs in AcCK2 vs AcT2 comparison group were found to target 14 miRNAs; among these DEcircRNAs, five DEcircRNAs can target two miRNAs, whereas another eight DEcircRNAs had only one target miRNA (**Fig. 4b**).

### 2.6 DEcircRNA-miRNA-mRNA regulatory network engaged in host response to *N. ceranae* infestation

Further investigation showed that 23 DEcircRNAs in AcCK1 vs AcT1 comparison group can target 18 miRNAs and further target 1111 mRNAs **(Supplementary Table S3)**. These target mRNAs were annotated to ten cellular component-related terms such as cell and membrane, eight molecular function-related terms such as catalytic activity and transporter activity, and 11 biological process-related terms such as cellular process and biological regulation **(Table S5)**. Additionally, these target mRNAs could also be annotated to 72 pathways including endocytosis, RNA transport, and ubiquitin-mediated proteolysis **(Table S6)**.

Additionally, 13 DEcircRNAs in AcCK2 vs AcT2 comparison groups can target 14 miRNAs and further target 1093 mRNAs **(Supplementary Table S4)**. These target mRNAs were engaged in ten cellular component-associated terms including organelle and cell, eight molecular function-associatedterms including binding and catalytic activity, and 11 biological process-associated terms including biological regulation and single-organism process **(Table S7)**. In addition, these target mRNAs could as well be engaged in 72 pathways including endocytosis, purine metabolism, RNA transport, protein processing of endoplasmic reticulum, and ubiquitin-mediated proteolysis, respectively **(Table S8)**.

Moreover, target mRNAs in both AcCK1 vs AcT1 and AcCK2 vs AcT2 comparison groups were involved in six cellular immune-related pathways, including endocytosis, lysosome, phagosome, ubiquitin mediated proteolysis, metabolism of xenobiotics by cytochrome P450, and insect hormone biosynthesis (**Table 7**); however, no target was annotated to any humoral immune pathway.

**Table 7.** DEcircRNAs engaged in Summary of cellular immune pathways enriched by mRNAs witin ceRNA networks.

| Pathway | Number of target mRNA | Number of target mRNA | Ko number |
| --- | --- | --- | --- |
|  | in AcCK1 vs AcT1 | in AcCK2 vs AcT2 |  |
| Endocytosis | 28 | 28 | ko04144 |
| Lysosome | 4 | 4 | ko04142 |
| Phagosome | 3 | 3 | ko04145 |
| Ubiquitin mediated proteolysis | 2 | 2 | ko04120 |
| Metabolism of xenobiotics by<br>cytochrome P450 | 1 | 1 | ko00980 |
| Insect hormone biosynthesis | 1 | 2 | ko00981 |

### 2.7 Investigation of DEcircRNAs’ protein-coding potential

Totally, 284 IRESs and 54 ORFs were identified from DEcircRNAs in AcCK1 vs AcT1 comparison group, respectively **(Supplementary Table S9-10)**. These ORFs were involved in two biological processes-associated terms, four molecular function-associated terms, and two cellular component-associated terms **(Supplementary Table S13)**. Additionally, these ORFs were engaged in eight pathways including endocytosis, other glycan degradation, ECM-receptor interaction, insulin resistance, other glycan degradation, FoxO signaling pathway, ribosome biogenesis in eukaryotes, ubiquitin-mediated proteolysis, and spliceosome **(Supplementary Table S14)**.

Comparatively, 164 IRES and 26 ORF were identified from DEcircRNAs in AcCK2 vs AcT2 comparison group **(Supplementary Table S11-12)**. These ORFs were enriched in two biological processes-related terms, four molecular function-related terms, and two cellular component-related terms **(Supplementary Table S15)**. In addition, these ORFs were involved in four pathways involving sphingolipid metabolism, ECM-receptor interaction, insulin resistance, and neuroactive ligand-receptor interaction **(Supplementary Table S16)**.

### 2.8 RT-qPCR validation of DEcircRNAs

Six DEcircRNAs were randomly selected for RT-qPCR validation, the result indicated that their expression trends were consistent with those in high-throughput sequencing data, which confirmed the reliability of transcriptime data used in this current work.

## 3. Discussion

In the present study, on basis of previously gained high-quality transcriptome data from *N. ceranae*-inoculated and un-inoculated midguts of *A. c. cerana* workers, the regulatory roles of circRNAs involved in *N. ceranae*-response of *A. c. cerana* workers were for the first time investigated. In workers’ midguts at 7 dpi and 10 dpi with *N. cerenae*, 10185 and 7405 circRNAs were resepectively identified, with a length distribution among 201∼800 nt (**Fig. 1c**), and annotated exon circRNA was the most abundant type (**Fig. 1b**). Similarly, we previously identified 6530 and 6289 circRNAs in un-inoculated workers’ midguts using the same bioinformatic approach, and found their length distribution was among 201∼800 nt and the most abundant type was also annotated exon circRNA (Chen et al., 2020). Further analysis showed that compared with un-inoculated groups, the number of circRNA enriched in each length or circularization type in *N. ceranae*-inoculated groups was totally higher than that in un-inoculated groups, implying that *A. c. cerana* workers may respond to *N. ceranae* infection by altering the quantity of circRNAs in differential length and circularization type. Additionally, 2266 circRNAs were shared by AcCK1, AcCK2, AcT1, and AcT2 groups, while the numbers of specific ones were 2618, 1917, 5717, and 3742, respectively (**Fig. 1a**), indicative of the developmental stage- and stress stage-specific expression of *A. c. cerana* circRNAs, which was similar to cricRNAs identified in other animals and plants (Hu et al., 2018; Liang et al., 2019). In view of that the information about circRNAs in Asian honey bees is very limited, circRNAs discovered here could well enrich the reservoir of *A. cerana* circRNAs. In addition, 83 and 52 DEcircRNAs were identified in AcCK1 vs AcT1 and AcCK2 vs AcT2 comparison groups, which suggested that the expression profile of circRNAs in host midguts was altered due to *N. ceranae* infection and these DEcircRNAs probably be involved in host response.

Accumulating evidence suggested that circRNAs can exert function through regulating transcription of source genes (Li et al., 2015; Zhang et al., 2013). *N. ceranae* has been proved to prolong the survival time of infected cells by inhibiting apoptosis of *A. mellifera* midgut epithelial cells to steal host material and energy for fungal proliferation (Kurze et al., 2018; Paris et al., 2018). Attionally, *N. ceranae* could also cause structural damage to midgut epithelial cells of *A. mellifera* (Panek et al., 2018). The Hippo signaling pathway plays crucial part in regulating cell proliferation and organ growth and regeneration (Tang et al., 2022). Emerging evidence indicated that Hippo signaling pathways played an pivotal role in regulation of immune defense of mammals and insects (Hong et al., 2018; Buchon et al., 2009). In this current work, it’s found that 78 source genes of 83 DEcircRNAs (novel_circ_008114) in AcCK1 vs AcT1 comparison group were engaged in the Hippo signaling pathway. It’s speculated that these DEcircRNAs may when detecting damage of midgut epithelial cell structure caused by *N. ceranae* infection, corresponding DEcircRNAs were employed by host to regulated source genes’ transcription, further modulating Hippo signaling pathway to facilitate cell renewal and regulate immune response.

Insect defends against pathogenic micriprganisms using cellular and humoral immune, and the former involves endocytosis, phagocytosis, enzymatic hydrolysis, melanization, and synthesis and release of antimicrobial peptides (Lavine and Strand. 2002). In honey bees, endocytosis is a main cellular immune pathways (Aronstein and Murray. 2010). Clathrin-mediated endocytosis is one of the most clearly studied endocytosis pathways, Kim et al. discovered that deletion of *FgEnd1* in *Fusarium graminearum* resulted in a significantly down-regulation of the endocytic marker FM4-64 and dicreaseof mycelium growth rate when compared with wild-type cells (Kim et al., 2009). Hodgson et al. found that progeny virions of *Autographa californica* multiple nucleopolyhedrovirus (AcMNPV) was significantly reduced when the early and late endosome marker genes Rab5 and Rab7 in *Drosophila* DL1 cells were disrupted by RNAi (Hodgson et al., 2019). Ubiquitin-mediated proteolysis is a classic cellular immune pathway, which involves E1 enzyme (ubiquitin activase), E2 enzyme (ubiquitin binding enzyme) and E3 enzyme (ubiquitin ligase); E3 can specifically recognize different substrates and then bind to E2 enzyme, which is finally recognized and degradated by protein enzyme body (Sutovsky, 2003). Here, it’s observed that two source genes of two DEcircRNA (novel_circ_005307; novel_circ_017023) in AcCK1 vs AcT1 comparison group were engaged in endocytosis, while one souce gene of DEcircRNA (novel_circ_016946) was involved in ubiquitin mediated proteolysis. This suggested that corresponding DEcircRNAs were likely to regulate the aforementioned two cellular immune pathways by regulation of sources genes’ transcription, and further participate in host *N. ceranae*-response. Intriguingly, in our previous work, it’s observed that source genes of DEcircRNAs in *A. m. ligustica* workers’ midguts during the *N. ceranae* infection were enriched in four cellular immune-related pathways including endocytosis, ubiquitin-mediated proteolysis, lysosome, phagosome (unpublished data). Collectively, these results indicated that both *A. c. cerana* and *A. m. ligustica* workers probably respond to *N. ceranae* invasion by regulating endocytosis and ubiquitin-mediated proteolysis through controling source genes’ transcription.

Increasing evidence suggested that circRNAs could regulated target gene expression via ceRNA network and further affect various biological processes such as immune response and development (Li et al., 2020; Han et al., 2020). Here, 23 and 13 DEcircRNAs in AcCK1 vs AcT1 and AcCK2 vs AcT2 comparison groups were predicted to target 18 and 14 miRNAs, further targeting 1111 and 1093 mRNAs, implying that these DEcircRNAs may exert function as ceRNAs during host response to *N. ceranae* infection. Further analysis indicated that target mRNAs in worker’s midgut at 7 dpi were associated with six cellular immune pathways including endocytosis, lysosome, phagosome, ubiquitin-mediated proteolysis, metabolism of xenobiotics by cytochrome P450, and insect hormone biosynthesis; whereas targets in worker’s midgut at 10 dpi were involved in five cellular immune-related pathways such as endocytosis, ubiquitin-mediated proteolysis, lysosomae, phagosome, and insect hormone biosynthesis. Interestingly, no target was detected to enrich in any humoral immune pathway. The results demonstrated that corresponding DEcircRNAs were likely to regulate host immune response to *N. ceranae* infection through ceRNA networks. miR-182 gene was found to be abundantly expressed in sensory organs and regulate development of retina and inner ear (Wei et al., 2015). In humans, Perilli et al. discovered overexpression of miR-182 could inhibit apoptosis and promote cell proliferation as well as colorectal cancer cell infection by altering tumor cell cycle dynamics and morphology (Perilli et al., 2019). Sun et al. revealed that miR-182 regulated RGS17 through two conserved sites within 3’UTR region, and ectopic expression of miR-182 conspicuously inhibited lung cancer cell proliferation and anchorage-independent cell growth (Sun et al., 2010). FOXO3a was previously identified as a direct target of miR-182-5p, and miR-182-5p played a inhibitory role in FOXO3a expression; activation of AKT/FOXO3a pathway can promote HCC proliferation and invasive ability, which resulted in higher death rates (Cao et al., 2018). Wu et al. discovered that miR-182-5p directly targets 3’ UTR of a tumor suppresser gene STARD13, which significantly relieved the inhibitory effect of decreased miR-182-5p on cell proliferation, migration, and invasion in lung adenocarcinoma (Wu et al., 2021). Here, it’s noted that miR-182 was targeted by two DEcircRNAs in AcCK1 vs AcT1 comparison group, including novel_circ_016924 (log_2_FC=17.25, *P*= 0.0020) and novel_circ_016946 (log_2_FC= 16.37, *P*=0.0077), indicating that these two DEcircRNAs may play a key part in cell apoptosis and cell proliferation by absorbing miR-182 in host response to *N. ceranae* infection.

Eukaryotic translation depends on the ribosomal scanning mechanism of the m7G cap structure (Haimov et al., 2015). Due to a lack of 5’ terminal and poly-A tail, circRNA was previously considered not to be able to translate protein. With the rapid development of next-generation transcriptome sequencing and ribosome profiling, some circRNAs were verified to translate into proteins or small peptides with biological functionvia an IRES-based manner (Wang et al., 2015; Pamudurti et al., 2017). Yang et al. reported that FBXW7-185aa, a protein encoded by circRNA FBXW7 (circ-FBXW7), plays a crucial role in glioma carcinogenesis and in patient clinical prognosis (Yang et al., 2018). After transfecting *Drosophila* S2 cells with artificially constructed circRNA including GFP gene containing IRES (Wang et al., 2015), Wang et al. detected that the constructed circRNA could successfully express green fluorescent protein (Wang and Wang, 2015). Weigelt et al. showed that overexpression of a protein-coding circRNA-circSfl could extend lifespan of insulin mutant Dorsophila (Weigelt et al., 2020). In this work, two DEcircRNAs in AcCK1 vs AcT1 comparison group (novel_circ_017023 and novel_circ_005307) were predicted to translate proteins relative to endocytic pathways, and one DEcircRNA in AcCK1 vs AcT1 comparison group (novel_circ_016946) was predicted to translate proteins associated with ubiquitin-mediated proteolysis pathway. This suggests that the above-mentioned three DEcircRNAs may be engaged in cellular immune response of *A. c. cerana* workers to *N. ceranae* invasion through translation of proteins relevant to endocytosis and ubiquitin-mediated proteolysis.

## 4. Materials and Methods

### 4.1 Honey bee and microsporidian

Three *A. c. cerana* colonies located in the teaching apiary of the College of Animal Sciences (College of Bee Science) in Fujian Agriculture and Forestry University were used for this study. Microscopic observation and PCR identification verified that these colonies were *N. ceranae*-free. No Varroa was observed before and during the whole experiment. RT-PCR was conducted to detect the prevalence of seven common bee viruses (DWV, KBV, ABPV, CBPV, IAPV, SBV, and BQCV) and two bee microsporidia (Nosema apis and *N. ceranae*) in the newly emergent workers with previously described specific primers (Chen et al., 2019a; Chen et al., 2008; Benjeddou et al., 2001; Genersch. 2005; Ribiere et al., 2002; Stoltz et al., 1995; Singh et al., 2010), and agarose gel electrophoresis (AGE) indicated that no bands specific for the aforementioned viruses and microsporidia were amplified (Chen et al., 2019a, Xing et al., 2021).

Foragers were collected from a *N. ceranae*-infected colony in an apiary in Fuzhou city, Fujian Province, China. *N. ceranae* spores were previously prepared using Percoll discontinuous centrifugation method by our group (Chen et al., 2019a).

### 4.2 Source of strand-specific cDNA library-based RNA-seq data

Midgut tissues of *A. c. cerana* workers at 7 dpi and 10 dpi with *N. ceranae* and corresponding un-inoculated workers’ midguts were previously prepared by our team (Xing et al., 2021). There were three biological replicates for each treatment or control group.

RNA isolation, cDNA library construction, deep sequencing, and data quality control were previously conducted (Xing et al., 2021). The constructed 12 cDNA libraries were sequenced on Illumina HiSeqTM 4000 platform (Illumina). Raw data are available in NCBI Short Read Archive database (http://www.ncbi.nlm.nih.gov/sra/) under BioProject number: PRJNA406998. In total, 1 809 736 786 raw reads were produced from RNA-seq, and 1 562 162 742 clean reads were gained after quality control; the mean Q20 of clean reads from control groups and treatment groups were 94.76% and 94.77%, respectively; the mapping ratio of clean reads to the reference genome of *A. cerana* were 75.78%(AcCK1), 55.01%(AcCK2), 78.13%(AcT1) and 44.19%(AcT2), respectively (Chen et al, 2020a; Fu et al., 2020).

The strand-specific cDNA library-based RNA-seq data with high-quality could be used for circRNA identification and investigation of DEcircRNAs in this work.

### 4.3 sRNA-seq data source

In another previous study, midgut tissues of *A. c. cerana* workers at 7 dpi and 10 dpi with *N. ceranae* and corresponding un-inoculated workers’ midguts were prepared by our team (Chen et al., 2019a). There were three biological replicas for each treatment or control group.

RNA extraction, cDNA library construction, sRNA-seq, and data quality control were previously performed (Chen et al., 2019a). The 12 constructed cDNA libraries were subjected to sequencing on Illumina MiSeqTM platform with the single-end strategy. A total of 127 523 419 raw reads were generated from sRNA-seq, and 122 104 443 clean reads were obtained after quality control; the Pearson correlation between every replica in each group was above 0.9619 (Chen et al., 2019a). The high-quality sRNA-seq data could be used for target prediction and regulatory network construction of DEcircRNAs in this study.

### 4.4 Bioinformatic prediction of circRNAs

CircRNAs in AcCK1 and AcCK2 groups had been identified in previous study (Chen et al., 2020a). In this work, circRNAs in AcT1 and AcT2 groups were identified following our previously described method (Chen et al., 2020a). Shortly, clean reads were mapped to the *A. cerana* reference genome (assembly ACSNU-2.0) using TopHat software (Kim et al., 2013), and 20 nt at both ends of unmapped reads were then extracted and independently mapped to the reference genome; the mapped anchor reads were submitted to find_circ software (Memczak et al., 2013) to perform circRNA identification following criteria: circRNA length < 100 kb, best qual A > 35 or best qual B > 35, anchor overlap ≤ 2, n uniq > 2, edit ≤ 2, n uniq > int (samples/2), and breakpoints = 1.

### 4.5 Identification of DEcircRNAs

The expression level of each circRNA was normalized to the mapped back-splicing junction reads per million (RPM) mapped reads value. Following the threshold |FC (fold change)| ≥ 2, *P* value < 0.05, and false discovery rate (FDR) ≤ 1, DEcircRNAs in AcCK1 vs AcT1 and AcCK2 vs AcT2 comparison groups were identified using DESeq software (Wang et al., 2010).

### 4.6 Analysis of source genes of DEcircRNAs

CircRNA could regulate the expression of source genes via interaction with RNA polymerase II, U1 ribonucleoprotein, and gene promoter (Li et al., 2015; Zhang et al., 2013). According to the method described by Chen et al (2020a), source genes of DEcircRNAs were predicted through mapping the anchor reads at both ends of DEcircRNA to the *A. cerana* reference genome (assembly ACSNU-2.0) using Bowtie software. GO term analysis of the circRNAs’ source genes was conducted with DAVID tool (http:// david.abcc.ncifcrf.gov/) (Huang et al., 2007) and the GO category was clarified using a two-sided Fisher’s exact test, while the FDR was calculated to correct the P value (Jung, 2014). KEGG pathway analysis was performed according to the annotation of the Kyoto Encyclopedia of Genes and Genomes (KEGG) database (http://www.genome.jp/kegg/) (Du et al., 2014).

### 4.7 Construction and investigation of DEcircRNA-miRNA and DEcircRNA-miRNA-mRNA regulatory networkers

In combination with previously identified miRNAs based on sRNA-seq data (Chen et al., 2019a), target miRNAs of DEcircRNAs were predicted on basis of the previously described protocol by Chen et al. (2020a). Briefly, according to the criteria of *P* ≤ 0.05 and free energy ≤ -35 kcal/mol, potential target miRNAs of DEcircRNAs were predicted using three software including mireap, miranda (v3.3a) and targetscan (version: 7.0), followed by construction of DEcircRNA-miRNA regulatory network; further, target mRNAs of DEcircRNAs-targeted miRNAs were predicted and then DEcircRNA-miRNA-mRNA regulatory network was constructed; the regulatory networks were visualized by Cytoscape software (Smoot et al., 2011) with the default parameter. GO term and KEGG pathway analyses of target mRNAs were further conducted employing the above-mentioned methods.

### 4.8 RT-PCR and Sanger sequencing of circRNAs

Three circRNAs (novel_circ_005123, novel_circ_007177, and novel_circ_015140) shared by AcT1 and AcT2 groups were randomly selected for molecular verification. Following our previously described method (Chen et al. 2020a), divergent primers for these circRNAs (Table 1) were designed using DNAMAN 8 software (Lynnon Biosoft, USA) and then synthesized by Shanghai Sangon biological co., Ltd. Total RNA of workers’ midguts at 7 dpi and 10 dpi with *N. ceranae* were respectively extracted with AxyPre RNA extraction kit (Axygen, China), and then digested with 3 U/mg RNase R (Geneseed, China) at 37°C for 15 min to remove linear RNA. Subsequently, the first-strand cDNAs were synthesized via reverse transcription with random primers. The PCR reaction system was 20 μL in volume containing 1 μL of template, 10 μL of Mixture (Yeasen, China), 1 μL upstream primers (10 μmol/L) and 1 μL downstream primers (10 μmol/L), and 7 μL ddH_2_O. The reaction was carried out on a T100 thermocycler (Bio-Rad, USA) following the conditions: 94°C for 5 min; followed by 36 cycles of 94°C for 40 s, an appropriate annealing temperature (according to the melting temperature of the primers) for 30 s, 72°C for 30 s; and 72°C for 5 min. The PCR products were detected on 1.5% AGE followed by TA cloning and Sanger sequencing.

### 4.9 RT-qPCR validation of DEcircRNAs

Four DEcircRNAs in AcCK1 vs AcT1 (novel_circ_010689, novel_circ_005734, novel_circ_016924, novel_circ_0176939) and two DEcircRNAs in AcCK2 vs AcT2 (novel_circ_008642, novel_circ_005927) were randomly selected for RT-qPCR. Divergent primer of these DEcircRNAs were designed and synthesized (**Table 8**). Total RNA of AcCK1, AcT1, AcCK2 and AcT2 groups were isolated and then subjected to reverse transcription, the resulting cDNA were used as templates for internal control (*actin*) and DEcircRNAs. RT-qPCR reaction system was 20 μL in volume containing upstream primers 1 μL (10.0 μmol/L), downstream primers 1 μL 10.0 μmol/L, cDNA 1μL, SYBR Green Dye 10 μL, DEPC H_2_O 7 μL. RT-qPCR were performed on an ABI Q3 Real-time PCR Detection System (Applied Biosystems, USA) under the following conditions: predenaturation step at 94°C for 5 min; 36 amplification cycles of denaturation at 94°C for 50 s, annealing at 60°C for 30 s, and elongation at 72°C for 1 min; a final elongation step at 72°C for 30 s. The data was calculated using 2^-△△Ct^ method (Livak and Schmittgen, 2001) and presented as relative expression levels from three parallel replicas and three biological replicas, followed by analysis and visualization using GraphPad Prism 6.0 software (GraphPad, USA).

**Table 8.**
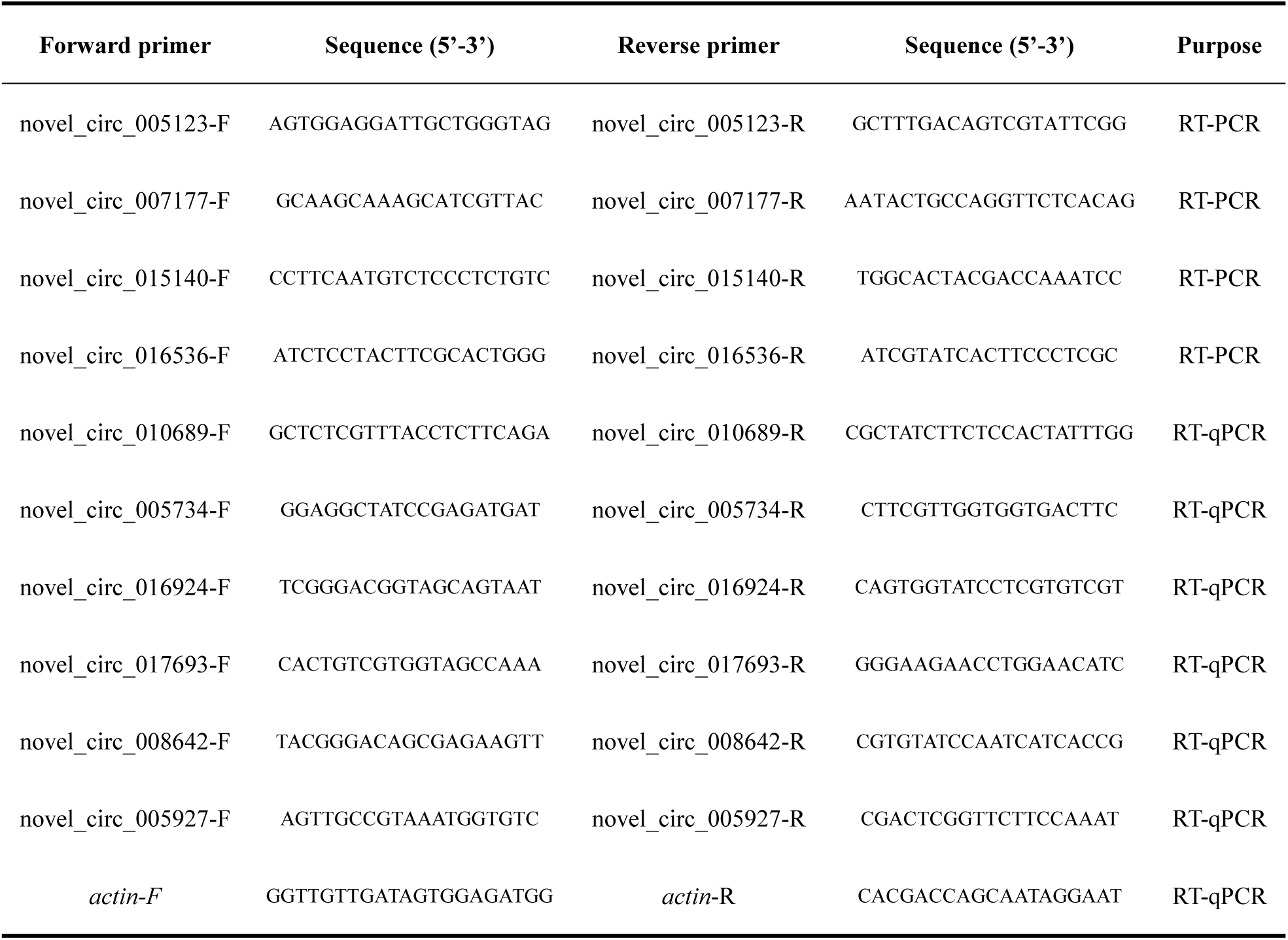
Primers for RT-PCR and RT-qPCR conducted in this work

## 4.10 Statistical analysis

Statistical analyses were conducted utilizing SPSS 16.0 (IBM, Armonk, NY, USA) and GraphPad Prism 6.0 (GraphPad, USA) software. Data were showed as mean ± standard deviation (SD). Statistical analysis was performed with independent-sample t test. Fisher’s exact test was conducted to filter the significant GO terms and KEGG pathways using R software 3.3.1 (R Development Core Team, https://www.r-project.org/). *P* < 0.05 was considered statistically significant.

## 5. Conclusions

In the present study, we for the first time investigated the expression profile and potential function of circRNA in *A. c. cerana* workers’ midguts responding to *N. ceranae* infection. The results indicated that the expression pattern of circRNAs was altered due to *N. ceranae* infection, DEcircRNAs may play regulatory roles in host immune response through regulation of transcription of source genes, as absorption of target miRNAs via ceRNA network, or translation of proteins. Our data offer a foundation for clarifying the mechanism underlying immune response of *A. c. cerana* workers to *N. ceranae* invasion and a novel insight into host-parasite interaction during bee nosemosis.

